# Automating 3DED data processing at eBIC

**DOI:** 10.64898/2026.07.29.741428

**Authors:** Marko D. Petrović, C. David Owen, David McDonagh, Daniel Hatton, Éilís C. Bragginton, Pedro Nunes, Adam D. Crawshaw, David G. Waterman

## Abstract

Three-dimensional electron diffraction (3DED) is an emerging and useful technique for solving molecular structures of small and biological macro-molecules from nanometre-sized crystals. We present our automated data processing workflow for 3DED datasets collected at Diamond Light Source’s electron Bio-Imaging Centre (eBIC). For this purpose, we developed a package called AutoED. The processing pipeline includes data collection, analysis of the beam position, metadata gathering, file conversion, and finally data processing using xia2 (which supports both DIALS and XDS). The processing results are captured in a summary report produced by AutoED. Our main goal is to reduce the workload of electron diffraction scientists, but also to enforce good standards already used in macromolecular crystallography (MX). All the collected 3DED datasets are automatically converted into NeXus data format which is considered a Gold Standard for MX. This standardized data format allows for all the relevant metadata about the experiment to be kept together with diffraction images. We also discuss the methods used in AutoED to determine the electron beam position on diffraction images.

## 1. Introduction

3DED is gaining attention as a technique for deriving structural information from microcrystalline samples. In part, this is driven by the increasing access to high-quality electron microscopes, for example through facilities such as the electron Bio-Imaging Centre (Clare *et al.*, 2017) at Diamond Light Source (DLS), improvements in detector technology and data collection strategy (Clabbers *et al.*, 2022) (Martynowycz *et al.*, 2022), improvements in sample preparation (Jones *et al.*, 2018) (Parkhurst *et al.*, 2023), and finally developments in the data analysis software to ensure that the measurements result in useful data. Automation of diffraction data processing was a valuable and necessary step in the continued development of single crystal X-ray diffraction. We are now at a point at which these developments can be deployed against electron diffraction.

### 1.1. Automation in X-ray Diffraction

3DED is following a path travelled by X-ray diffraction, and many other scientific disciplines, with data quality and quantity increasing with improvements in instrumentation: rapidly the volume of data becomes too great for one scientist to process manually in a timely manner. Since the development of high throughput techniques to support structural genomics, and in particular the Protein Structure Initiative (Montelione, 2012) automated processing has become routine at X-ray diffraction facilities for structural biology, to the point where this is considered an essential part of the beamline system (Monaco *et al.*, 2013) (Winter & McAuley, 2011). These automated processing systems benefit from the investment of effort in developing automation-friendly data processing packages - most conspicuously XDS (Kabsch, 2010) and DIALS (Winter *et al.*, 2018) as well as automated “pipelines” to simplify integration of the software into a beamline environment (Winter *et al.*, 2013; Vonrhein *et al.*, 2011).

These investments in data processing software, and the pipelines to run them, simplify the application of those tools to more complex data processing challenges, including new pipelines to approach massively-multi-crystal data processing such as multiplex (Gildea *et al.*, 2022). In short, the result of these developments is that a scientist’s first interaction with a dataset can be inspection of an electron density map of a site of interest, highlighting the presence or absence of a ligand. This can be followed by careful reprocessing if required, but importantly it allows the experiments to be guided by the biology.

### 1.2. Challenges relating to Electron Diffraction

Given the existence of tried-and-tested automated pipelines for X-ray diffraction data processing, why can these not simply be adopted immediately for 3DED? The differences between X-ray and electron single crystal diffraction are well documented and adaptations to software such as DIALS have been made to optimise results for 3DED (Clabbers *et al.*, 2018; Vypritskaia *et al.*, 2025). For automated data-processing the greatest impediment remains inaccurate or incomplete experimental metadata. Unfortunately, the typical file formats used in electron microscopy are not designed for diffraction, and do not transmit the required metadata faithfully. Reliable and reproducible capture of experimental metadata has been a long-standing focus point for synchrotron X-ray diffraction. NXmx has been adopted as essentially complete and a high-fidelity expression of the experimental data and metadata, ensuring the data are useful to future users and can be revisited if and when software and practices improve. With adaptions this format can be used for electron diffraction (Waterman *et al.*, 2023). An important point for automation is that the required metadata must be accurate, but must not impose an excessive burden on instrument scientists or the experimentalist. Practically, if the data can be automatically and satisfactorily processed at acquisition time with metadata provided by the experimental setup, then the metadata are demonstrably sufficient.

A challenge that arises in particular for 3DED is that the beam centre is not in general known prior to data collection, and may be subject to experiment-to-experiment variation. This causes problems with indexing the diffraction data, i.e. determining the unit cell dimensions and internal symmetry of the crystal system. A robust and automated method of determining the beam centre is necessary to avoid a significant amount of manual “busy” work on the part of the scientist, and is critical for the implementation of a fully-automated pipeline.

Problems with indexing are compounded by noisy results from spot-finding and the typically limited rotation range of 3DED experiments due to sample mounting on electron diffraction grids. Above a certain angle, either extinction, alignment or the grid mount obscuring the beam eliminates the diffraction. While instrumentation and workflows have improved significantly over recent years (Aragon *et al.*, 2024), the challenges inherent to 3DED data collection mean that some datasets are likely to be compromised due to experimental issues rather than crystal quality alone. For example, misalignment between the rotation axis and sample leads to a horizontal offset at larger rotation angles, potentially moving the sample out of the beam. This will give rise to datasets where only the middle of the scan contains useful data with the edges presenting minimal diffraction. Faced with multiple data sets of varying quality, automated data processing becomes not just a convenience for the experimentalist but an essential initial filter to select the promising datasets. At synchrotrons, tools have been developed to be run in parallel with data acquisition, including live processing, symmetry determination from many small wedges of data (Gildea & Winter, 2018) and automated combination and analysis of sample isomorphism, orientation, and overall reciprocal space coverage. As many of the same issues are present, these tools are equally applicable to electron diffraction data.

### 1.3. Motivation

Available electron diffraction instrumentation currently allows rapid acquisition of data from many samples on a grid, operating in an automated mode, translating from one crystal position to the next. It would be easy to overwhelm the experimentalist if relying on manual data processing alone. Importantly, offering automated processing does not preclude the scientist from returning to datasets later for more careful investigation of tricky cases. One clear motivation for offering automated processing is to allow the scientist to focus on the biological question, such as “how does my ligand bind?”, rather than procedural challenges relating to format conversion and manual recording of the beam centre.

“Live” feedback, on a similar timescale to the measurement of a single scan, is valuable. Early warning that the samples are not isomorphous can be useful as an indicator that additional care or a modified collection strategy will be needed to get complete data, and allows the experiment strategy to be optimised to make the best use of microscope time.

To bring electron diffraction further into the mainstream we must reduce barriers to non-specialists. Irrespective of the philosophical considerations, building experience on how to process diffraction data takes time, which can be in short supply. If, therefore, we have the capability to offer good quality automated processing we have an obligation to do so, to ensure that the maximum benefit may be drawn from the use of the samples and instrumentation. Several automated pipelines for 3DED data processing have been described in recent years. Scipion-ED (Bengtsson *et al.*, 2022) extends the Scipion cryo-EM workflow framework to electron diffraction, providing a graphical user interface (GUI) for processing (based on DIALS as the backend). AutoLEI (Wang *et al.*, 2026) works with XDS as the backend engine, and provides both real-time and offline batch processing of many datasets. Its GUI provides reports to assess the data quality of processed results. Automation has also been extended beyond data reduction: AutoMicroED (Powell *et al.*, 2021) runs from raw images through to structure solution using SHELX for small molecules and Phaser for proteins, while Instamatic-solve (Luo *et al.*, 2025) couples XDS and SHELXT for structure solution.

## 2. Results and Discussion

### 2.1. AutoED package

Crystal structure determination is a multi-step process, from sample preparation, acquiring high-quality images, and detecting diffraction spots to integration and determining atomic positions. Some steps of this procedure are well-defined in that they accept a standardised input and produce a standardised output. An example of this is the processing of NeXus files using xia2 (spot finding, indexing, integration, scaling, etc.). xia2 can accept a variety of input formats and provides standardised, predictable output (scaled and integrated `MTZ files`). Solving the structure after the spot intensities are already determined is also addressed by dedicated software such as SHELX or Phaser. The remaining challenge for full automation, therefore, lies in data acquisition and metadata handling prior to the xia2 processing step. This is still an open challenge due to the variety of file formats used to save diffraction images and the lack of a coherent effort by equipment (microscope and detector) manufacturers to adopt a uniform standard. The automation of this part depends heavily on the current experimental setup in each lab, and, given their heterogeneity, it is very hard to standardize across labs.

This paper addresses the issue of automating the meta-data collection so that xia2 processing becomes possible. To automate the processing of ED data collected at eBIC, we developed a Python package called AutoED. It is an open source command-line tool and is available through the Python Package Index (PyPI).^1^ AutoED is a wrapper around existing diffraction software, such as xia2 and DIALS, and focuses primarily on metadata handling that enables diffraction integration. Our goal is to remove the need for electron diffraction scientists to manually extract metadata from microscope log files and images, and embed this metadata in files that can be processed with xia2 or DIALS. The first step in that direction was the extension of the Nexgen package (Waterman *et al.*, 2023) to 3DED data, allowing the conversion of several electron diffraction data formats into NeXus (e.g. the plain HDF5 SINGLA detector format). The NeXus format is chosen as the best way to standardise input for xia2 because it is already considered a Gold standard in MX and defines a hierarchical structure for all relevant ED metadata.

Another open problem addressed by AutoED was the absence of critical metadata, namely the electron beam position. By default, the DECTRIS SINGLA detector does not record the position of the electron beam on diffraction images. This piece of information is crucial for successful data processing, because without it, no experimental model can be constructed. Extracting the beam position by visually inspecting diffraction images (e.g. in DIALS image viewer) is not a difficult task, but repeating that for thousands of datasets very quickly becomes impractical.

### 2.2. AutoED workflow

The basic AutoED workflow implemented at eBIC is presented in Fig. 1. Experimental datasets are copied from the Glacios microscope to the DLS file system. AutoED runs as a daemon on one of the DLS servers with full access to this file system. The daemon can start and stop watchdog processes that monitor specific directories. In addition to the diffraction images, the microscope PC generates and transfers a small JSON file containing relevant metadata about the experiment.

**Figure 1.**
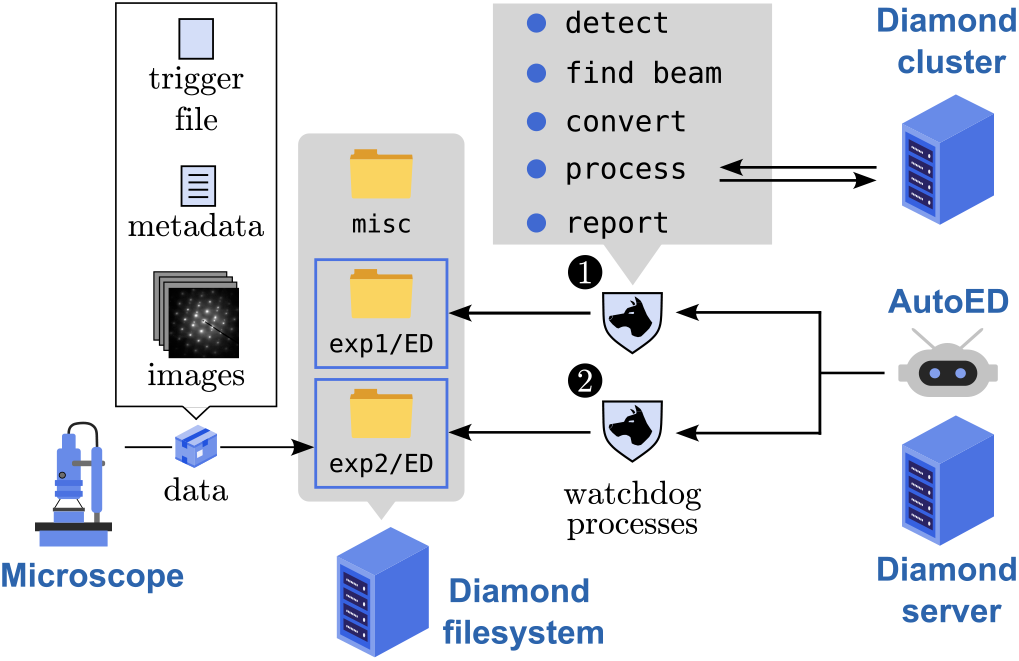
Schematics of the AutoED workflow: The data generated by the eBIC microscope is transferred to the Diamond Light Source (DLS) file system, where it is detected and processed by an AutoED watchdog process. After detection, the data is analysed to determine the beam position and converted to NeXus format (with the new beam position included in the metadata). Next, the data is processed on the DLS cluster, and processing statistics are included in a report file.

ED datasets are relatively large files (several gigabytes in size). Therefore, the time of detecting a new file might not coincide with a dataset being entirely copied from the microscope. To prevent incomplete data from being processed, the microscope PC sends a trigger file after copying a dataset to signal to AutoED that the dataset is ready for processing. After detecting a new dataset, a watchdog process will:

1. Analyze the dataset and determine the position of the beam center (which we will discuss in the sub-section below).
2. Convert the output DECTRIS SINGLA data file into the MX Gold standard NeXus format using the Nexgen package. Both the JSON metadata and the new beam position are embedded in the NeXus file. At this step, data and metadata are bound together at the point of collection, ensuring that metadata is not lost or separated from the data at a later stage.
3. Process the new NeXus files using xia2 with several choices of pipeline.
4. Generate a summary report for a better overview of the processing statistics. For each successfully processed dataset, xia2 generates a detailed report. However, when processing many datasets simultaneously, a higher-level overview is needed to quickly assess the quality across all processed datasets.

AutoED operates in two modes: a local mode, where datasets are processed sequentially one at a time, and a remote mode, where processing is submitted to the DLS cluster. To support this workflow, we established several conventions that are worth noting for anyone wishing to adapt AutoED to a different experimental setup.

- We require specific naming conventions; for example, image data and metadata must share the same base name to be interpreted as being part of the same dataset.
- Each watchdog process monitors only one directory at a time, but monitoring is recursive within that directory.
- No two watchdog processes can monitor the same directory, preventing the same dataset from being processed multiple times.
- As a control mechanism, processing is triggered by an empty file generated by the microscope PC after each dataset is transferred.

Processing of each dataset is done using a variety of pipelines. A pipeline consists of one or more (DIALS or xia2) commands run on a particular dataset. The pipelines are defined by template scripts that use placeholder variables whose values are substituted at runtime. By relying on template files, we allow the user to define their own processing steps. This allows the processing to be modular. The user can selectively enable and disable specific pipelines based on their current needs and resource availability, add new pipelines, or remove obsolete ones. Another advantage of this approach is that any changes to the command-line interface of the underlying processing tools do not affect AutoED itself, since the commands are defined in configuration files rather than hard-coded into the package.

This modularity, combined with xia2’s ability to run both DIALS and XDS to process data, allows for performing comparison runs with different parameter setups and quickly selecting the optimal ones. Furthermore, the user is not limited to running only xia2 or DIALS; they can include their own processing scripts or any other command-line tools on their system. Finally, for each dataset, the pipeline template is used to generate a bash processing script that accompanies the original data, so that at any point, the record of the commands used to process the data is preserved. The user can rerun these bash scripts manually, supporting the reproducibility aspect of FAIR principles. At its core, the processing part of AutoED is a template-based bash script generator.

During the processing, it is important to keep track of the processing statistics. xia2 already generates report files for individual datasets; however, the user needs an overview of how processing is progressing across all collected datasets. The purpose of AutoED reports is to provide a rapid means of distinguishing high-from low-quality datasets. The user should be able to discard datasets that could not be processed, and rapidly select datasets based on specific criteria (high number of indexed spots, good CC1/2 coefficient, high completeness). Additionally, when processing fails or produces unexpected results, it is essential to quickly identify the cause of the failure (e.g., incorrect beam position, incorrect mask, noise at a specific resolution). A clear, interactive summary of processing statistics can significantly speed up this process. The AutoED report is an interactive HTML page containing a table with one row per processed dataset.

Any of the previously listed responsibilities of a watchdog process (finding beam position, processing the dataset, generating reports) was designed to spawn a separate process, thus not affecting the calling watchdog script. For each dataset, the watchdog is creating a sequence of pipeline processes, each of which updates HTML report. The hierarchy presented in Fig. 2 reflects the composition of the AutoED package. There is a central daemon class, a watchdog class, a dataset class, a pipeline class, and a report generator class. They are all a part of the processing chain.

**Figure 2.**
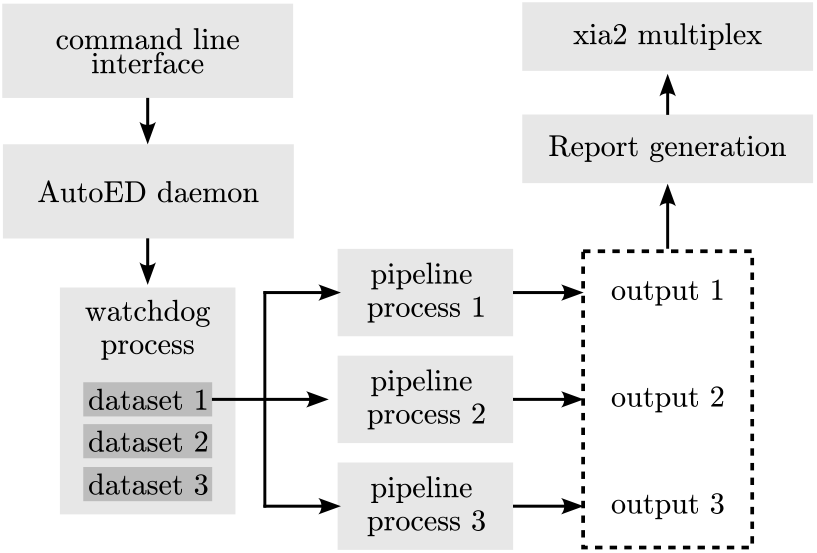
The internal structure of the AutoED package: the central part of the package is the AutoED daemon that can start and stop watchdog processes, which can begin individual pipeline processes. A watchdog class relies on a dataset class (which defines what a dataset is and what operations can be performed on it). A datasets class calls a pipeline class to process the dataset, and after processing, a report generator object parses the xia2 results and creates the report document. Additional object runs xia2 multiplex on the successfully processed datasets.

### 2.3. Methods to determine electron beam position on diffraction images

The most critical step in automatic metadata collection is determining the electron beam position on diffraction images. Several approaches can be taken to solve this problem: developing a machine-learning model (e.g., a convolutional neural network), fitting a profile to the beam position (e.g., a Gaussian), or processing image pixels to compute the most likely position of the beam. We opted for the third approach, as it requires no training data, is straightforward to interpret, and is computationally efficient. Without the use of an energy-filter, the amount of inelastic scattering in ED often produces a visible region of high intensity (a halo) around the beam position. This can be a problem for determining the intensity of low-value Miller indices (close to the direct beam). On the other hand, this scattering can be used to our advantage: the halo is usually symmetrically distributed around the direct beam. As we will show below, this symmetry is central to our approach.

Determining the beam position can be decomposed into two independent problems: finding the position in the x and y directions separately. The first step is to project the image data along these two directions, reducing a two-dimensional problem to two independent one-dimensional problems. This projection also significantly reduces the amount of data we operate on: for example, a 1000×1000 pixel image yields two projected arrays of 1000 values each, rather than a single array of 1,000,000 pixels, making our procedure computationally efficient. The methods we developed rely on these projected profiles as their primary input, but differ in how they extract the beam position from them. We call these methods the maximum pixel method, the inversion method, and the midpoint method, and we briefly outline each one below.

The SINGLA detector used at eBIC consists of two active panels separated by a horizontal gap (Fig. 3(a)). This gap contains no detector sensors, so the corresponding pixels are always masked. In some images, the beam is positioned within this gap region, rendering the beam effectively invisible in the image—we only see an indirect halo around it. In images where the beam is directly visible (outside the gap), determining its position is straightforward: one simply identifies the highest-intensity pixel.

**Figure 3.**
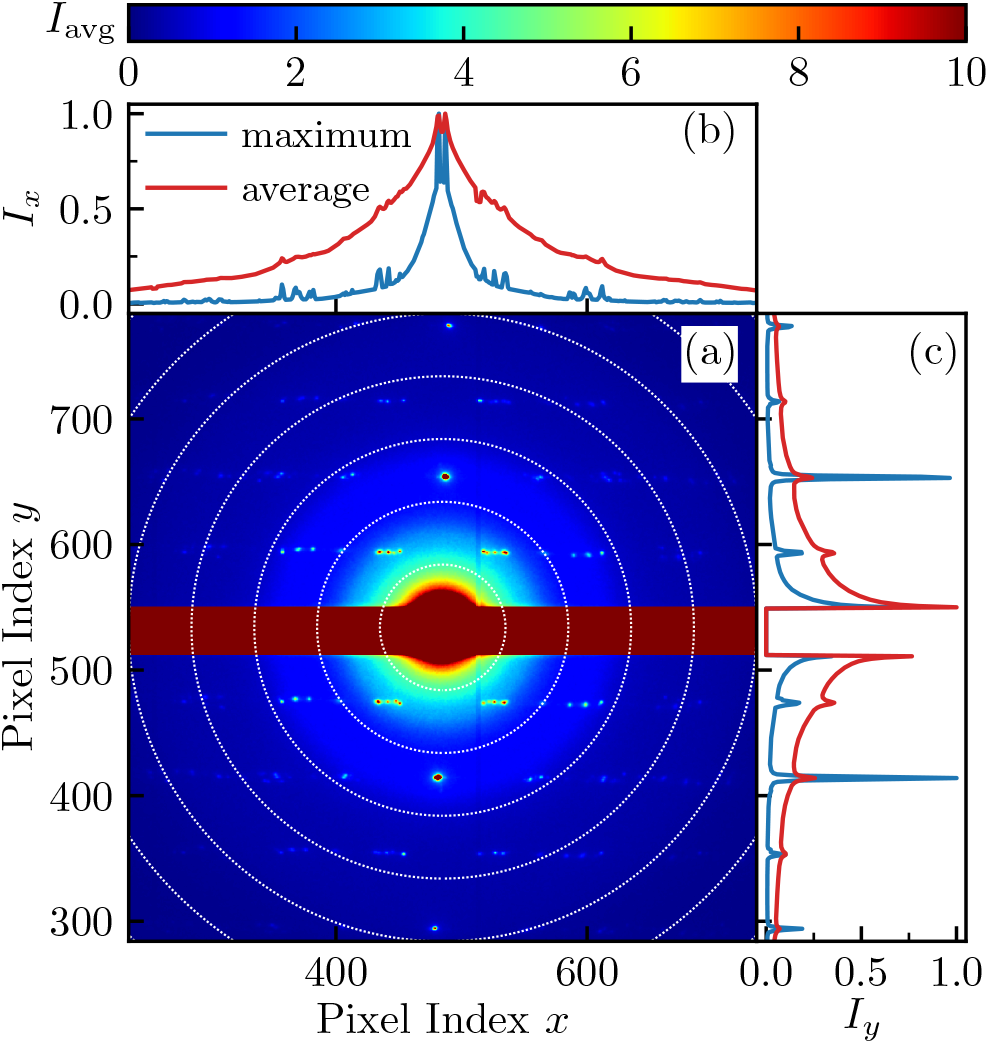
Computing beam center from projected profiles. (a) Average rotation diffraction image from a SINGLA dataset centered around the electron beam. (b) Normalized maximal and average image projection from (a) along the *x* direction. (c) Normalized maximal and average projection from (a) along the *y* direction.

The first step in determining the beam position is therefore to classify each image as either one where the beam is hidden (in the gap) or visible (outside the gap). Based on this classification, AutoED chooses the appropriate method: the maximum pixel method for a visible beam, or the midpoint method for a hidden beam.

Of these methods, the maximum pixel method is the simplest to describe. In this method, the beam position corresponds to the position of the pixel with maximum intensity in the projected profile (e.g. see the blue curve in Fig. 3(b)). Using only the maximum pixel can be problematic in some cases, for example when there are unmasked bad pixels or when some of the diffracted spots have the highest intensity in the entire image. To address this, before picking the highest pixel, we narrow the search area to the region most likely to contain the direct beam.

When the beam is hidden in the detector gap, the maximum pixel method is no longer applicable, and we must rely on the indirect halo around the beam. The second method we developed, the inversion method, is designed for this case. It is based on the properties of Friedel pairs and the assumption that the diffraction pattern is centrosymmetric around the direct beam. As shown in Fig. 3(a), looking at an average diffraction image, one can see pairs of spots with similar intensity clustered around the direct beam, which acts as a center of inversion. In principle, the simplest way to determine the beam position would be to connect these Friedel pairs with vectors and compute their midpoints. In our approach, since we work with projected profiles, these vectors are reduced to one dimension (*x* or *y*).

The main assumption of the inversion method is that the beam position acts as a center of inversion of the projected profile. If the original profile in Fig. 4(a) (gray curve, copied from Fig. 3(c)) is inverted at this point, the Friedel peaks in the original profile would coincide with those in the inverted profile. The goal is then to find this center of inversion. This is achieved by systematically inverting the projected profile at every pixel position and computing the overlap with the original curve. This overlap can be quantified as the integral of the product of the original and inverted profiles. One such inverted profile is shown as a dashed blue curve in Fig. 4(a), computed very close to the actual beam position, resulting in a clearly visible overlap. Fig. 4(a) also shows the overlap integral for every pixel along the *y* direction. The beam position is identified as the maximum of this overlap curve.

**Figure 4.**
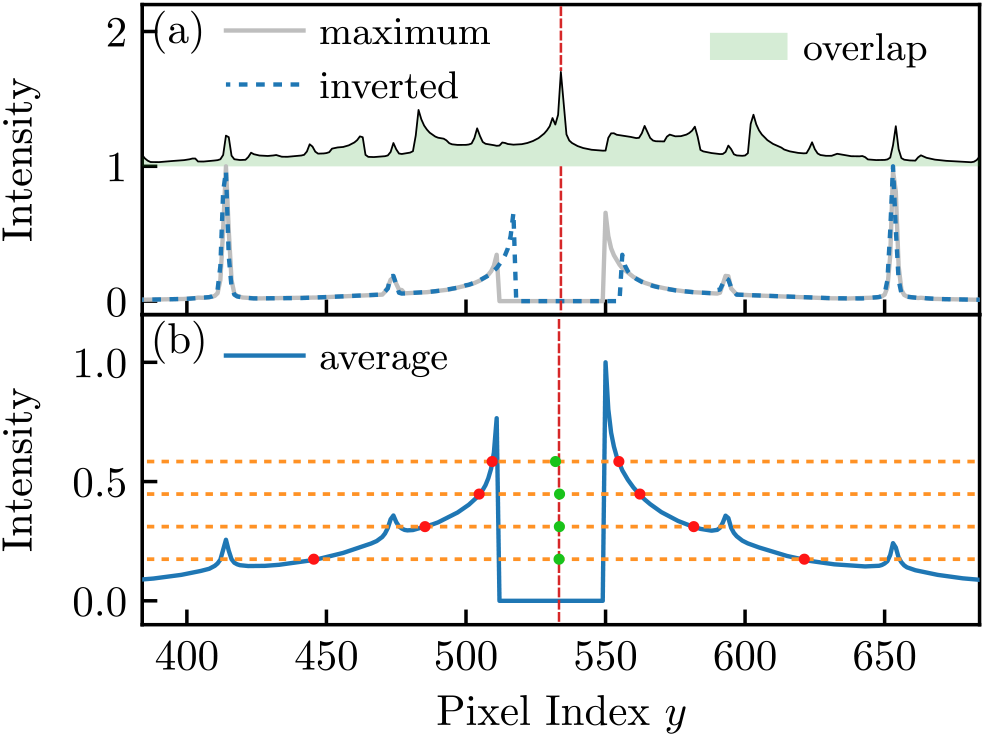
(a) Computing the beam position using the inversion method. The normalized maximal projection (gray curve) is inverted for every pixel index, and we compute the overlap function (shaded green area). The overlap function is scaled and shifted vertically for better visibility. The dashed blue curve shows the inverted profile at the computed beam position (marked with vertical dashed red line). (b) The beam position computed using the midpoint method. The normalized average projected profile (blue curve) is intersected with horizontal lines. We compute the intersection points (red dots) and the midpoints (green dots). The beam position is calculated as the average midpoint position.

Finally, the midpoint method assumes that the distribution of the scattered electrons around the direct beam is symmetric. In this approach, a set of horizontal lines (Fig. 4(b)) is drawn through the projected average profile, and the intersections of these lines with the projected profile are computed (red dots in Fig. 4(b)). The midpoint is then calculated for each pair of intersection points (green dots in Fig. 4(b)), and the beam position corresponds to the mean of these midpoints. Compared with the previous two methods, the midpoint method has a key advantage: it is applicable regardless of whether the beam is visible or hidden, and does not rely on the presence of Friedel pairs, which may not always be clearly visible in noisy datasets.

The methods described here rely on general properties of electron diffraction images: the strong intensity of the direct beam, the symmetric distribution of scattered electrons around it, and the inversion symmetry of Friedel pairs. These properties have been used in previous works to determine the beam position; see (Sikorova *et al.*, 2026) and references therein for an overview of existing methods.

### 2.4. Sensitivity of the processed result to beam position and oscillation angle

Whether a dataset is processed depends on the accuracy of determining the beam position and the rotation increment angle during the automatic data processing. The beam center for a single dataset is determined from the average rotation diffraction image. At the same time, the angle increment (oscillation) is read from the microscope by multiplying the goniometer rotation speed with the detector framerate. Figure 5 shows how varying beam position from its actual value influences the processing results, namely the percentage of spots indexed and the CC_1/2_ coefficient. We picked three Lysozyme datasets with 99%, 81%, and 77% indexed spots (from now on, datasets 1, 2, and 3, respectively), representing datasets with different crystal/diffraction quality. The presented results give an estimate of how precise the beam position needs to be for a dataset to be processed successfully and with a high CC_1/2_ coefficient.

**Figure 5.**
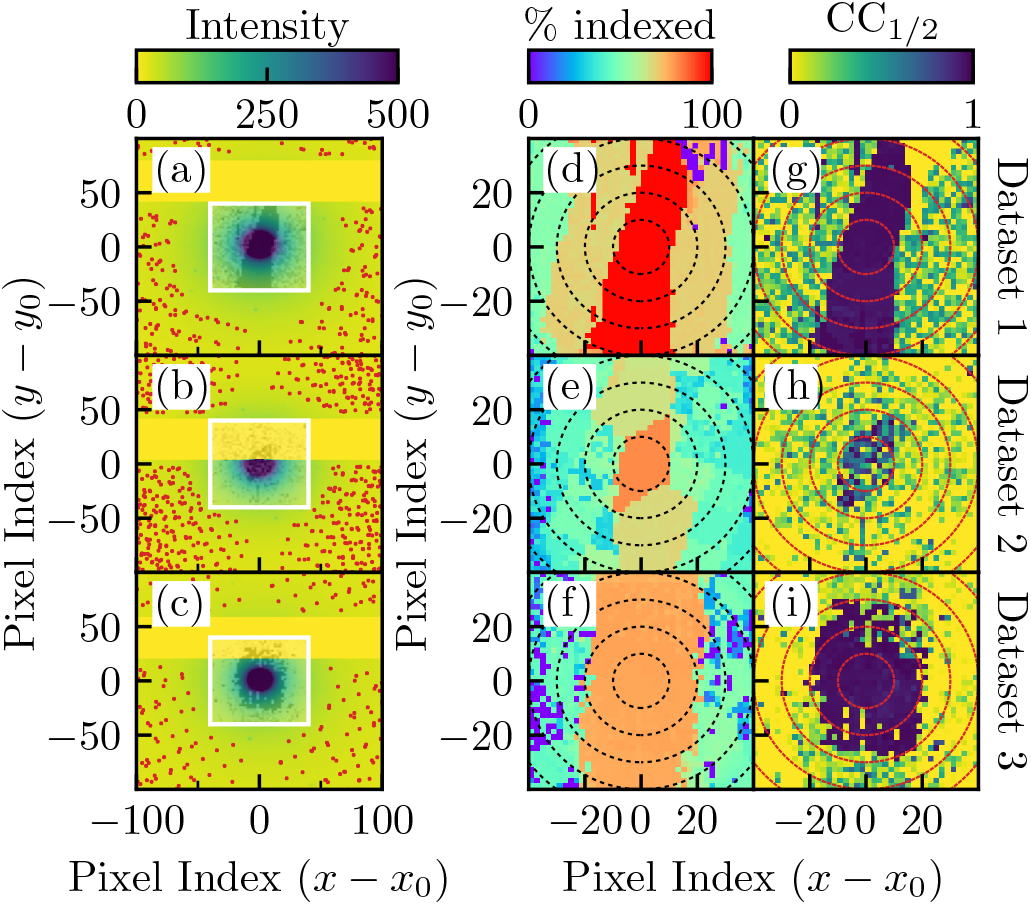
The sensitivity of the processed results to the position of the beam center. (a), (b), (c) Average rotation diffraction image with indexed spots (red dots) for three Lysozyme datasets, respectively. All images are centered around the actual beam position (*x*_0_, *y*_0_). (d), (e), (f) The percentage of indexed spots as a function of the imposed beam center for the corresponding three datasets, respectively. The imposed beam center varies in the *±* 40-pixel range around the actual beam center in both directions. The resolution of the scan in both directions is two pixels. (g), (h), (i) the CC_1/2_ coefficient as a function of the changing beam center for the corresponding three datasets. The beam center varies in the same way as in (d), (e), and (f). The white rectangles in (a), (b), and (c) show the region over which we varied the beam position, with the corresponding CC_1/2_ scans shown as a transparent overlay. The dashed circles in (d)-(i) are eye guides marking the distance from the actual beam position in ten-pixel radius increments.

For all three datasets, the results are unaffected if the beam position is within the ten-pixel radius of its actual position. On the other hand, rapid oscillations of the CC_1/2_ coefficient in Fig. 5(d) (even within the ten-pixel radius from the actual beam position) show that results change from dataset to dataset and that the quality of processed results can change drastically even for slight deviations in the computed beam position. All of the scan images in Fig. 5(d) - 5(i) show there is a vertical area in which the results do not change with the moving beam. This area possibly coincides with the rotation axis projected on the average diffraction image.

Another variable that potentially influences the quality of the processed results is the rotation angle increment Δ*ϕ*. As shown in Figure 6, the CC_1/2_ coefficient, the percentage of indexed spots, and the unit cell volume highly depend on small changes in the rotation angle increment. This parameter determines where to place the observed spots in the reciprocal space, and can therefore highly influence the determination of the unit cell parameters. As a consequence, the unit cell volume can change, or some spots might not be recognized as belonging to the reciprocal lattice of a given crystal. The most drastic effect is observed for the first dataset, where going above the default value of Δ*ϕ* = 0.5^°^ disrupts the processing of the entire dataset. The sensitivity of the processed results to changes in these two important parameters highlights the necessity to determine them as accurately and precisely as possible. The processing software under the control of xia2 will do some refinement of the beam position, so there is a small room for error; however, the precision of the rotation angle is crucial for building the correct crystal model.

**Figure 6.**
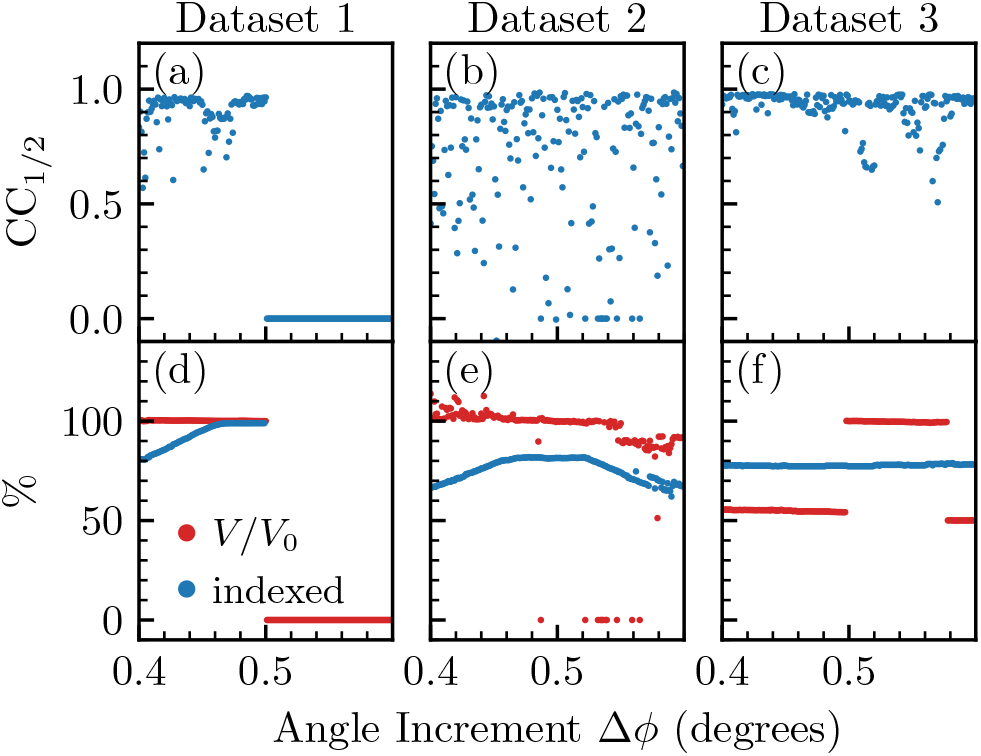
The sensitivity of processed results to rotation angle increment Δ*ϕ*. (a), (b), (c) The sensitivity of the CC_1/2_ coefficient for the three datasets considered in Fig. 5. The default angle increment Δ*ϕ* obtained from the microscope for all three datasets is 0.5 degrees. (d), (e), (f) The corresponding percentage of indexed spots (blue dots) and the relative change in the unit cell volume (red dots) for the three datasets from (a), (b), and (c), respectively. *V*_0_ is the unit cell volume at the default angle of Δ*ϕ* = 0.5 degrees. The angle increment is varied with 0.001 degree resolution. Data that failed to process is plotted at CC_1/2_ = 0 in the first row or at the zero percentage in the second row.

## 3. Conclusion

We developed AutoED, an open source command-line Python package for automated processing of electron diffraction data. AutoED builds on existing diffraction packages, such as xia2 and DIALS, and uses Nexgen to automatically embed all relevant experimental metadata into the NeXus format prior to processing. The package automatically determines the beam position from diffraction images using three methods: the maximum pixel method, the inversion method, and the midpoint method. We showed that the accuracy of the determined beam position, as well as the correct oscillation angle, are critical for successful data processing, and that even small deviations from the true values can significantly affect the quality of the processed results. The package also generates an interactive HTML summary report, allowing researchers to quickly filter datasets and identify processing failures. AutoED is currently in active use at eBIC, where it processes datasets on a timescale comparable to data acquisition, providing researchers with near real-time feedback during experiments. By automating metadata handling, format conversion, and processing, AutoED contributes to the broader goal of making 3DED data collection and processing compliant with FAIR principles, ensuring that all relevant metadata is preserved alongside the diffraction data. Although AutoED was developed for a specific experimental setup at eBIC, the conventions and configuration-driven design mean that adapting it to other setups requires only changes to configuration files rather than modifications to the core package.

Despite the progress described here, there is still work to be done to achieve full end-to-end automation of the 3DED pipeline. The remaining manual step is data acquisition, in particular the selection of high-quality crystals, which currently requires a skilled operator to visually inspect and select candidates. A logical extension of this work would be to automate this step through a package capable of automatic crystal classification and selection, with an interface to navigate to and position the electron beam on pre-selected crystals. With the growing development of Machine Learning models for computer vision, the problems of detecting crystals in images (classification), determining their position (object detection), and determining their morphology, that is, the precise region to illuminate with the electron beam (segmentation), become addressable. Several efforts are already underway to develop such automated procedures (Yonekura *et al.*, 2021; Eremin *et al.*, 2025). Given that selecting high-quality crystals is still a skill gained through experience, the expectation is that a faster rate of data acquisition and processing will compensate by allowing high-quality crystals to be filtered from a larger dataset rather than manually selected by an expert. In this way, AutoED represents a step toward bringing 3DED automation to the level already established in macromolecular X-ray crystallography.

## 4. Acknowledgements

This work was supported by Ada Lovelace Centre and BBSRC grant number BB/Y009991/1.

## 5. Supplementary Materials

### 5.1. Calibration of Rotation Speed

Successful and predictable automated processing of standard samples is an excellent sign that the metadata being provided to the software is correct. These metadata will include parameters such as sample to detector distance, oscillation width, and beam centre. Oscillation width is dependent on stage rotation speed, as collection for a defined period of time with a defined detector frame rate, this defines the oscillation per frame. Both stage rotation and detector distance speed need to be calibrated on a per microscope basis and should be monitored over time. As demonstrated in Fig. 6 in the main text, errors in oscillation per frame can directly impact success or failure of the indexing and refinement steps of diffraction data processing. A method for initial calibration of stage rotation speed of a Thermo Scientific Glacios microscope using SerialEM is provided below. To demonstrate the importance of calibrations such as these we collected a triclinic lysozyme dataset with a 130° sweep range divided into 0.5° oscillations. Detector distance can be calibrated using standards such as evaporated carbon, which provide a known powder diffraction pattern.

This serialEM script can be used to generate a csv file containing *time elapsed* and *angle*, from which change in angle over time can be plotted and a rotation speed in °/s determined.

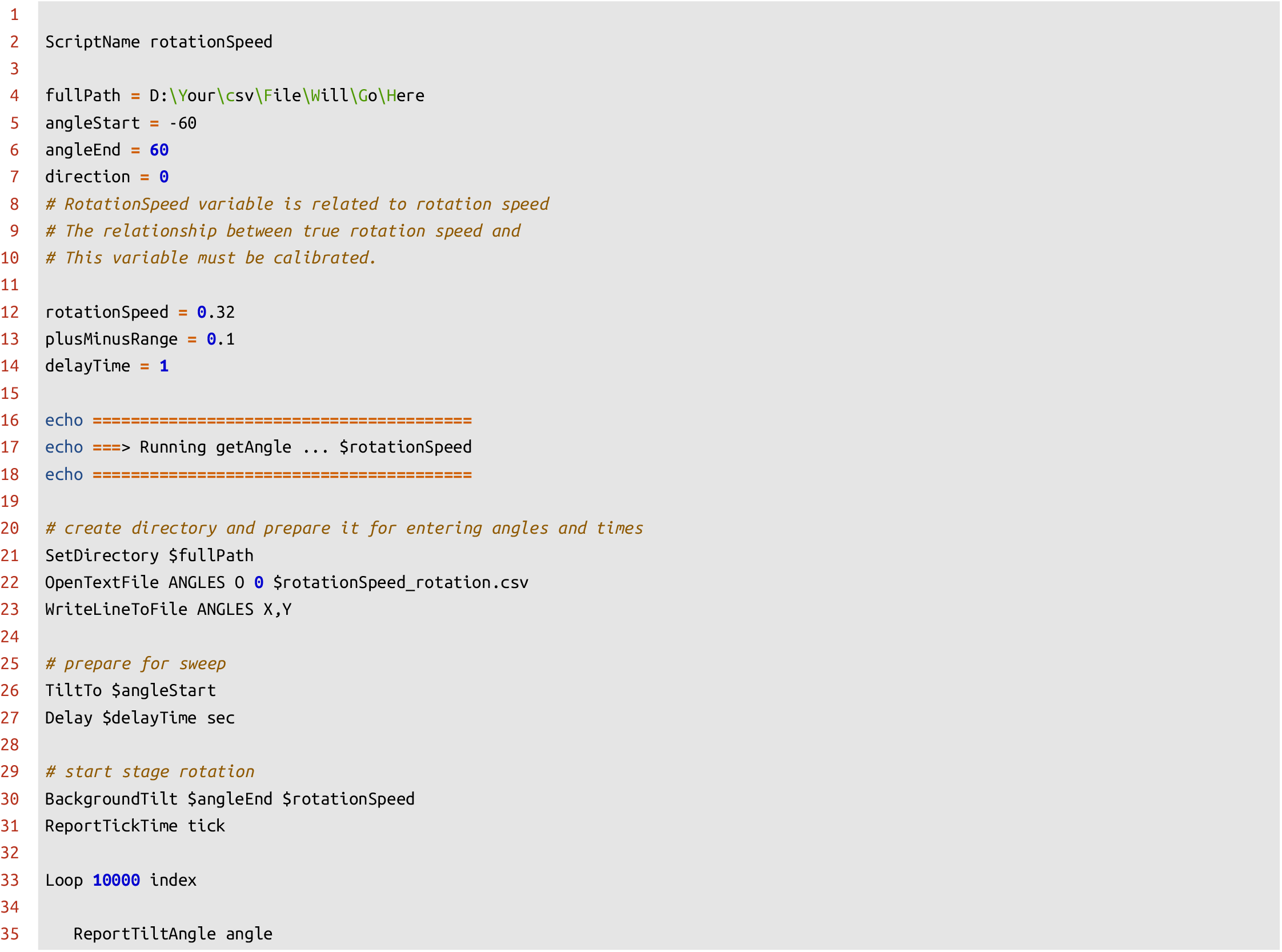
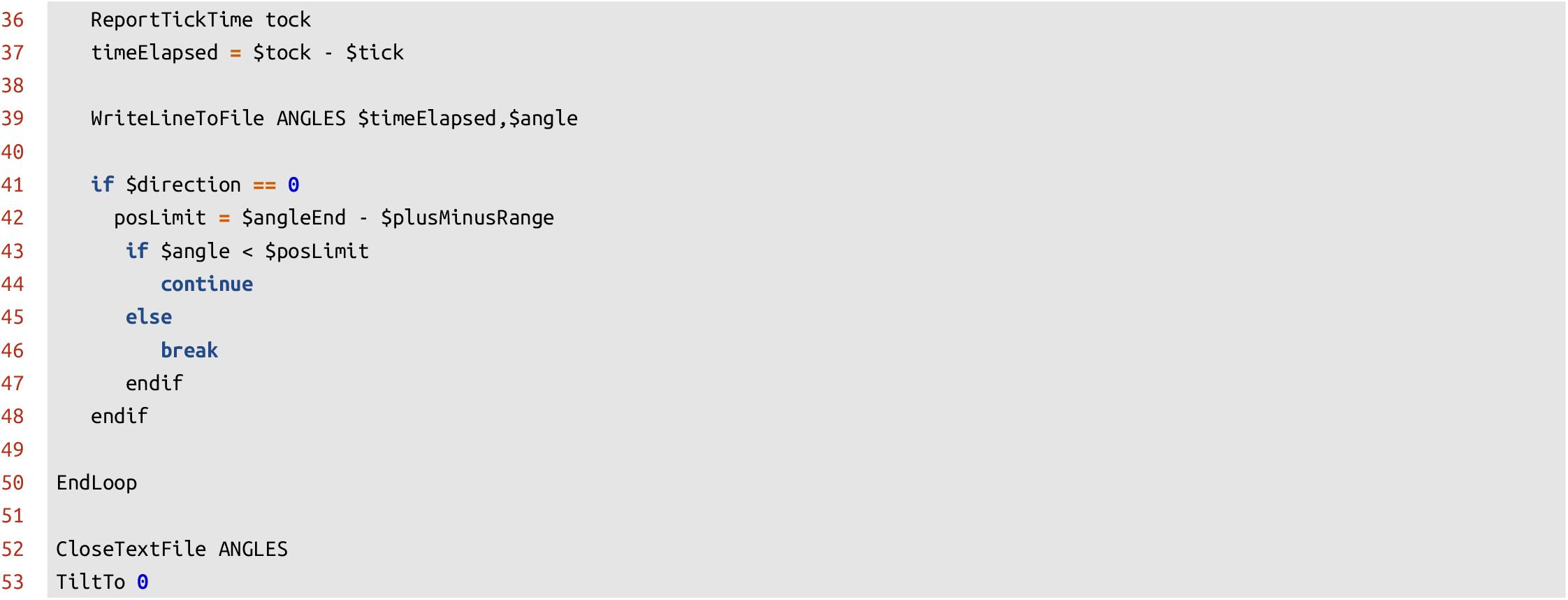

We provide an example python script that can be used to determine the true rotation speed from data generated by the above serialEM script.

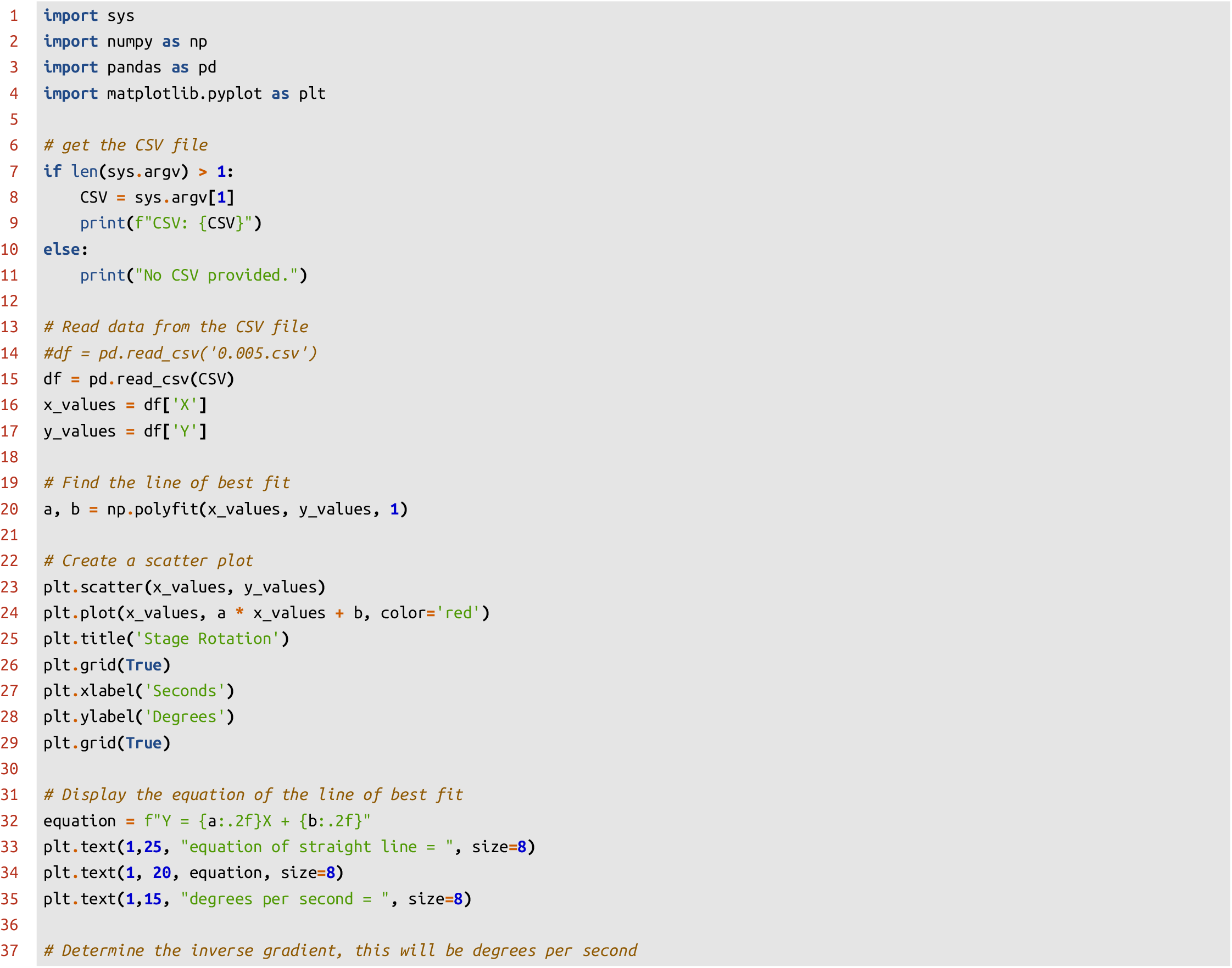
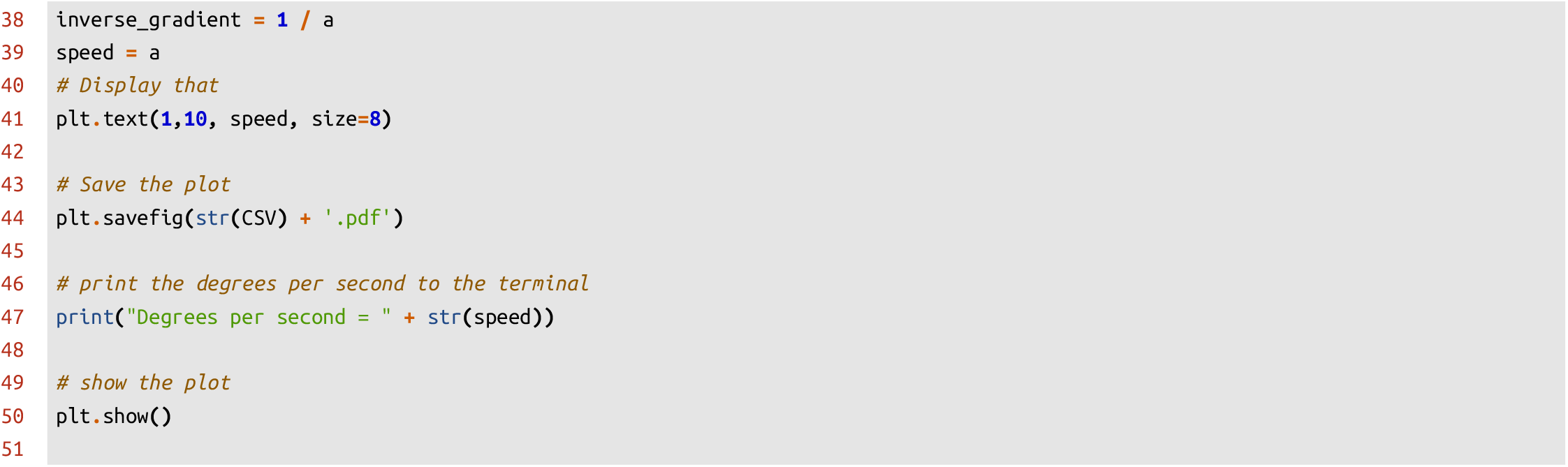

Please take care to inspect the stage rotation plots as they will indicate the presence or absence of “wind-up” and “wind-down” time, during which the stage is accelerating or decelerating. One should not collect data during these times as the oscillation angle per frame will not be constant (see Fig. 7). The presence of “wind-up” and “wind-down” time becomes more apparent when faster stage rotation speeds are attempted. They can be accounted for introducing a delay between starting stage rotation and the commencement of diffraction data collection (see Fig. 8). Care should be taken to ensure that variable stage rotation speed does not influence the calibration of stage rotation speed. These rotation speed calibration experiments should be repeated at sensible intervals, such as after manual stage interventions, or when indicated by poor data processing results from well understood standard samples.

**Figure 7.**
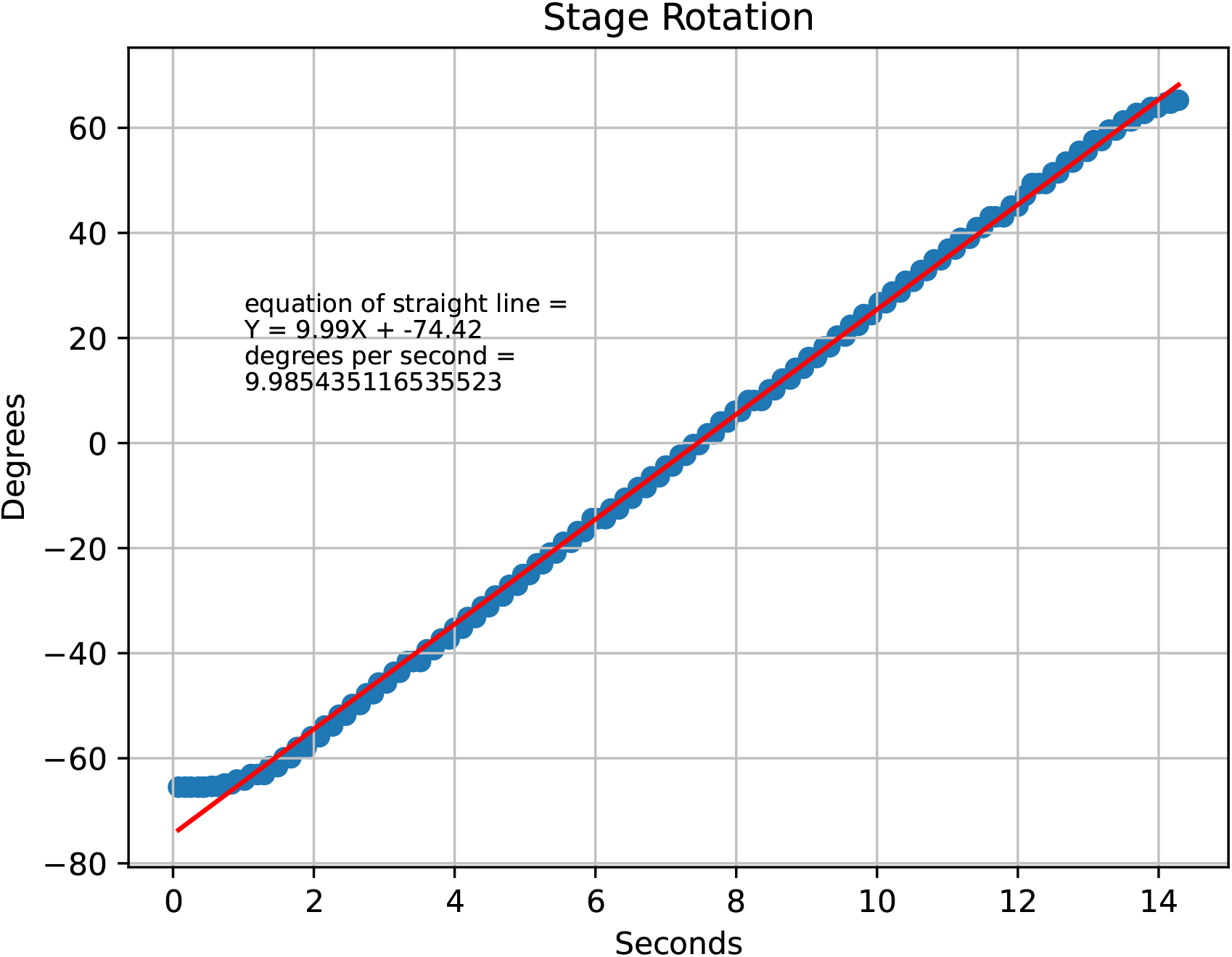
Change in stage rotation over time with a 0.25 second delay before commencement of angle recording.

**Figure 8.**
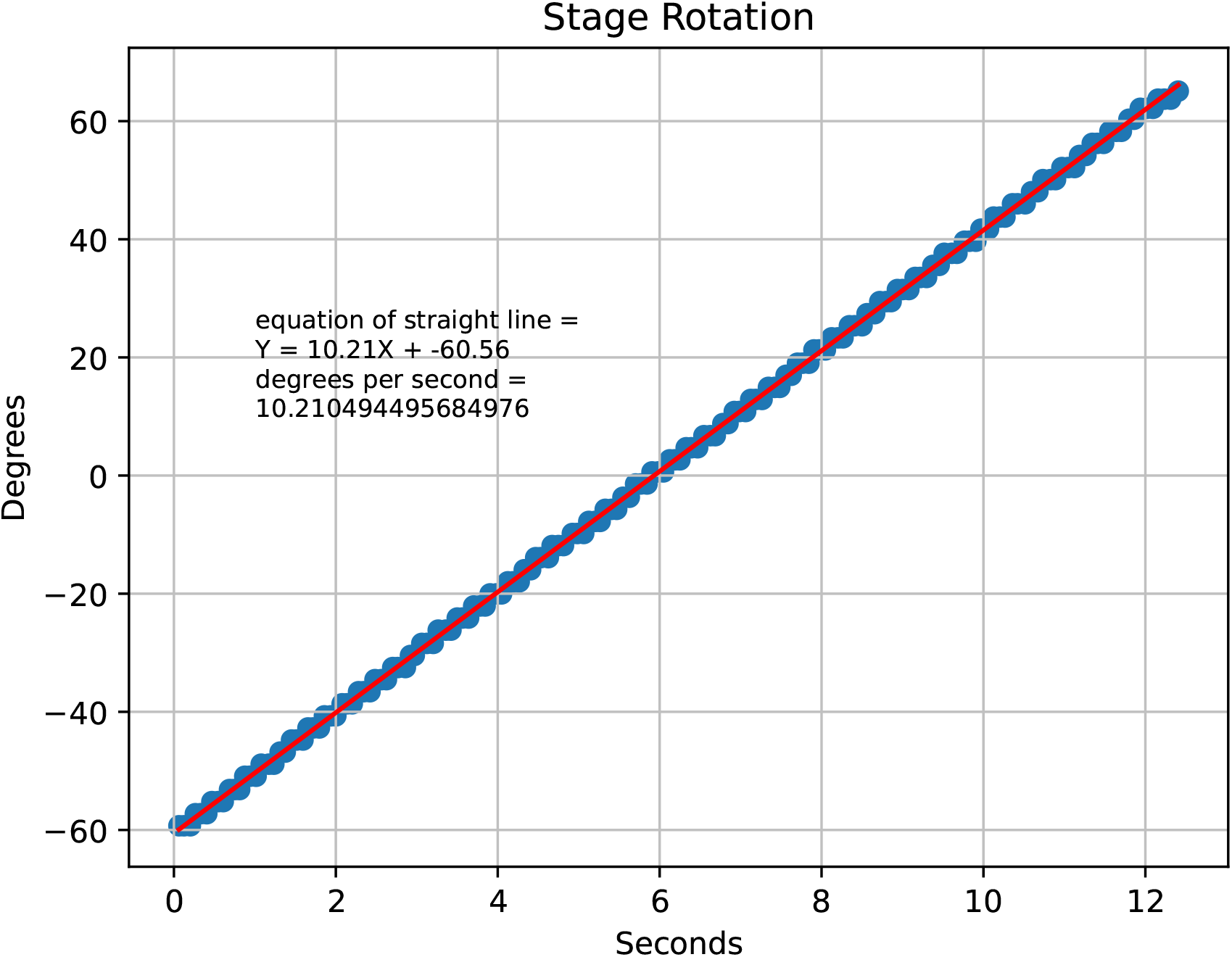
Change in stage rotation over time with a 2 second delay before commencement of angle recording.

## Footnotes

1 https://pypi.org/project/autoed/

